# RAD52 underlies the synthetic-lethal relationship between BRCA1/2 and 53BP1 deficiencies and DNA polymerase theta loss

**DOI:** 10.1101/2022.03.20.485027

**Authors:** Katarzyna Starowicz, George Ronson, Elizabeth Anthony, Lucy Clarke, Alexander J. Garvin, Andrew D Beggs, Celina M Whalley, Matthew Edmonds, James Beesley, Joanna R Morris

**Affiliations:** Birmingham Centre for Genome Biology and Institute of Cancer and Genomic Sciences, College of al and Dental Schools, University of Birmingham, B15 2TT, UK; Genomics Birmingham, College of Medical and Dental Schools, University of Birmingham, B15 2TT, UK; University of Leicester, UK; Insight Medical Writing Ltd, Danebrook Court, Oxford Office Village, Kidlington, Oxfordshire, OX5 1LQ, UK

**Keywords:** Homologous Recombination, BRCA1, 53BP1, RAD52, BRCA2, DNA polymerase theta, TMEJ

## Abstract

Cells lacking several DNA repair proteins, including those promoting homologous recombination (HR), are sensitive to polymerase theta (Polθ) repression^1–4^. Polθ drives theta-mediated end joining (TMEJ) and suppresses HR but what mediates its synthetic lethal relationships is unclear. Here we examine murine *Brca1^C61G/C61G^ 53bp1^-/-^*cells and find they are largely HR proficient by using RNF168 and RAD52. They exhibit no more TMEJ than *53bp1^-/-^* cells yet are more sensitive to targeting of Polθ. We find that RAD52 recruitment to damaged chromatin is increased following Polθ depletion. RAD52 accumulation and cellular sensitivity to Polθ repression can be curbed by the RAD51-binding regions of BARD1 and BRCA2, and sensitivity of BRCA1/2 depleted cells to Polθ repression is suppressed by RAD52 inhibition*. 53bp1^-/-^* cells exhibit a smaller increase in RAD52 recruitment following Polθ repression and also become resistant to Polθ repression following RAD52 inhibition. Thus, RAD52 mediates sensitivity to targeting Polθ in these contexts.

## Main

Inheritance of mutations in the <u>br</u>east <u>ca</u>ncer predisposition *BRCA1* or *BRCA2* genes carries an elevated risk of breast and ovarian cancer. These genes are central to the process of homologous recombination (HR) repair, and current targeted therapies, such as poly(ADP-ribose) polymerase inhibitors (PARPi) and platinum-based agents, aim to exploit HR vulnerability^5^. Cells lacking BRCA1 or BRCA2 proteins are sensitive to targeting of other repair pathways, including the activities of DNA polymerase theta (Polθ), encoded by the *POLQ* gene^2, 3, 6^. The recent development of Polθ inhibitors show they are able to kill *BRCA* deficient cells and cancers^7, 8^ and cancer cells lacking components of the 53BP1-Shieldin complex^4, 8^, which may be lost in *BRCA1* deficient cancers^9^. Inhibition of Polθ has been suggested as an alternative or adjunct to PARPi or platinum-based agents for the treatment of HR-deficient tumours^7, 8, 10^.

Polθ has several roles; it mediates translesion synthesis^11^ and Theta-mediated end-joining (TMEJ) (also referred to as alternative end-joining or micro-homology mediated end-joining), and suppresses HR^2, 3, 6^. In TMEJ Polθ acts to synapse overlapping microhomology in ssDNA followed by extension from annealed ends ^12^. The helicase domain of the protein displaces Replication Protein A (RPA) bound to ssDNA to enable the annealing of short sequences of homology, using its polymerase domain to fill in the gap. TMEJ is suppressed by BRCA2^13–16^ and RAD52 ^17^, and Polθ promotes the survival of cells deficient in several repair pathways ^1–4, 18^. However, while proteins involved in HR directly suppress TMEJ, it is not clear if the reverse is true. Both repair mechanisms share a substrate of resected, single-stranded DNA (ssDNA) 3’ ends, and may compete with one another ^3^; additionally, targeting Polθ results in increased accumulations of RAD51 into foci, which in turn contribute to toxicity^2, 4^. Thus, the underlying cause of toxicity of Polθ repression in several clinically-relevant contexts is not clear.

Here we examine the HR competency and Polθ dependency of murine cells bearing the hypomorphic *Brca1^C61G^* allele^19, 20^ and lacking *53bp1^21^*. We find that the C61G-BRCA1:BARD1 heterodimer promotes HR repair in combination with RNF168 and RAD52. *Brca1^C61G/C61G^ 53bp1^-/-^* cells are sensitive to Polθ depletion. Furthermore, BRCA1:BARD1, BRCA2 and Polθ restrain inappropriate engagement of RAD52. Importantly, depletion or inhibition of RAD52 suppresses several deleterious impacts of Polθ targeting on *Brca1^C61G/C61G^ 53bp1^-/-^* cells, including correcting RAD51 kinetics, chromosome aberrations and cell viability. These data highlight RAD52 as a possible biomarker and potential source of Polθ inhibitor resistance.

Crossing *Brca1^+/C61G^* with *53bp1^-/-^* mice resulted in *Brca1^C61G/C61G^ 53bp1^-/-^* pups born at the expected Mendelian ratios. Using anti-Brca1 antibodies we precipitated proteins from *Brca1^+/+^ 53bp1^-/-^* and *Brca1^C61G/C61G^ 53bp1^-/-^* mouse embryonic fibroblasts (MEFs). Mass spectrometry analysis identified BRCA1 peptides including the residue encoded by codon 61 (Supplemental Fig. 1a), confirming expression of the mutant protein. Expression levels of BRCA1 and its co-dependent binding partner, BARD1, were 20 (± 5)% of those seen in cells with WT-BRCA1; foci of the mutant BRCA1-BARD1 heterodimer in irradiated cells were fainter, and the number of proximity-linked ligation foci of co-located BRCA1-BARD1 was at approximately 30 (± 10)% of that seen in WT cells (Fig. 1a and Supplemental Fig. 1b-d). These data show that *Brca1^C61G/C61G^ 53bp1^-/-^* cells express a reduced level of the BRCA1-BARD1 heterodimer.

**Fig. 1.**
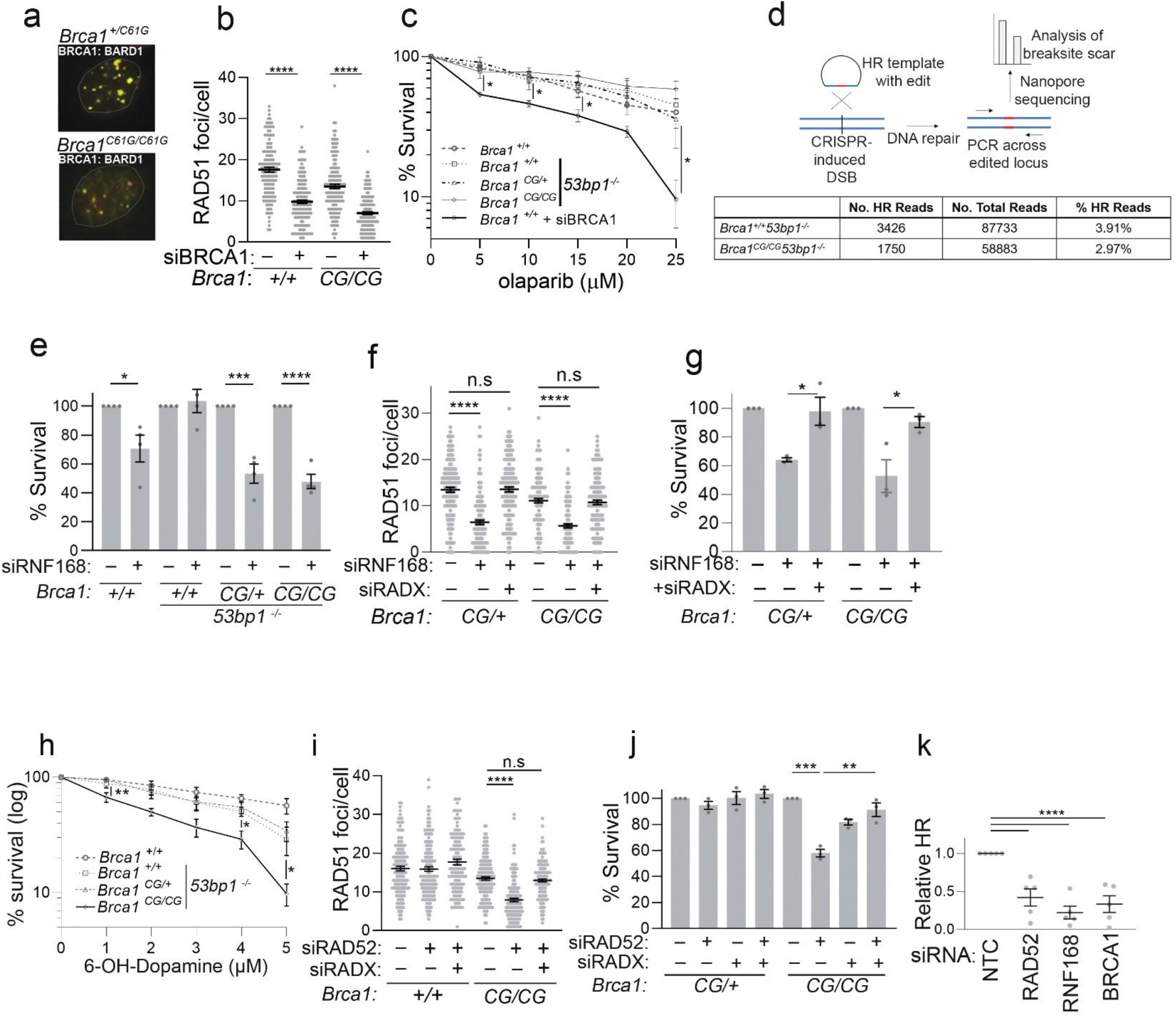
*Brca1^C61G/C61G^ 53bp1^-/-^* cells rely on non-canonical support mechanisms for RAD51 foci, HR and survival. **a**. Representative images of BRCA1 (red) and BARD1 (green), with yellow showing co-location, in the genotypes shown (all are *53bp1-/-*) exposed to 2 Gy irradiation. **b**. Quantification of RAD51 foci 3 hours after 2 Gy IR exposure, in EdU-positive MEFs of the genotypes shown and treated with NTC siRNA (-) or siRNA targeting BRCA1 (+). n = 150 cells from 3 biological replicates, per condition. Data are mean ± s.e.m. **c**. Colony survival following 16 h treatment with olparib measured in MEFs of the genotypes shown, or in WT cells treated with BRCA1 siRNA transfection, n=3, data are mean ± s.e.m. **d**. Illustration of the HR assay (top). The table below shows the % of reads containing the 4 bp inclusion and the total number of reads. **e**. Colony survival in MEFs of the genotypes shown treated with treated with NTC siRNA (-) or siRNA targeting RNF168 (+). n= 4, 4 biological repeats Data are mean ± s.e.m. **f.** Quantification of RAD51 foci 3 hours after 2 Gy IR exposure, in EdU-positive MEFs of the genotypes shown (all are *53bp1^-/-^*) and which were treated with NTC siRNA (-) or siRNA to RNF168, RADX or both (+). n = 150 cells from 3 biological replicates, per condition. Data are mean ± s.e.m. **g**. Colony survival in MEFs of the genotypes shown (all are *53bp1^-/-^*) treated with NTC siRNA (-) or siRNA to RNF168, RADX or both (+). n= 4, 4 biological repeats Data are mean ± s.e.m. **h.** Colony survival in MEFs of the genotypes shown (all are *53bp1^-/-^*) treated with the RAD52 inhibitor 6-OH-L-Dopamine. n= 4, 4 biological repeats. Data are mean ± s.e.m. **i**. Quantification of RAD51 foci 3 hours after 2 Gy IR exposure, in EdU-positive MEFs of the genotypes shown (all are *53bp1^-/-^*) and which were treated with NTC siRNA (-) or siRNA to RAD52, RADX or both (+). *n* = 150 cells from 3 biological replicates, per condition. Data are mean ± s.e.m. **j.** Colony survival in MEFs of the genotypes shown (all are *53bp1^-/-^*) treated with NTC siRNA (-) or siRNA to RAD52, RADX or both (+), n= 3, 3 biological repeats Data are mean ± s.e.m. **k.** Relative PCR product intensity of HR-specific *Vs* locus specific product in *Brca1^C61G/C61G^ 53bp1^-/-^* cells treated with NTC siRNA (-) or siRNA to RAD52, RNF168 and BRCA1. n=3 biological replicates, data are mean ± s.e.m.

We explored features of HR in *Brca1^C61G/C61G^ 53bp1^-/-^* cells. We noted that they exhibited both BRCA1-dependent RAD51 foci following irradiation (IR) or cisplatin treatment and BRCA1-dependent survival, did not show elevated levels of radial chromosomes after hydroxyurea treatment, and were resistant to poly(ADP-Ribose) polymerase (PARP1/2) inhibition by olaparib (Fig. 1b & c, Supplemental Fig. 1e-h). These phenotypes are commonly used to define HR competency. To quantitate HR further, we examined the ability of these cells to incorporate a homologous template into DNA cut by CRISPR/Cas9 at a specific site. Nanopore sequencing of the edited locus from *Brca1^+/+^ 53bp1^-/-^* cells, showed 3.91% of reads contained the HR product, versus 2.97% from *Brca1^C61G/C61G^ 53bp1^-/-^* cells, a deficit of 24% (Fig. 1d). Estimating HR using semi-quantitative PCR with primers specific for the HR outcome yielded a deficit of 48% (Supplemental Fig. 1i & j). Therefore, despite *Brca1^C61G/C61G^ 53bp1^-/-^* cells exhibiting some features of competent HR, the outcome of HR repair is sub-optimal.

We therefore explored mechanisms reported to support HR in cells with reduced or absent HR proteins. RNF168 promotes BRCA1-BARD1 recruitment^22^ and RAD51 loading through PALB2 interaction ^23^, ^24^. In *Brca1^C61G/C61G^ 53bp1^-/-^* cells we found that RNF168 depletion had a limited impact on BARD1 foci but substantially reduced the numbers of RAD51 foci formed following IR treatment, and reduced cell survival in otherwise untreated *Brca1^C61G/C61G^ 53bp1^-/-^* and *Brca1^C61G/+^ 53bp1^-/-^* cells (Supplemental Fig. 2a & b, Fig. 1e & f). Further, co-depletion with the ssDNA binding competitor of RAD51, RADX^25, 26^, restored both RAD51 foci numbers observed after IR treatment and cell survival to near control levels (Fig. 1f & g).

We then investigated a potential role for RAD52 as it promotes several recombination-mediated repair and replication mechanisms following DNA resection^27^. We found that depletion of RAD52 or treatment with a small molecule inhibitor of RAD52, (6-OH-D)^28^, decreased the viability of *Brca1^C61G/C61G^ 53bp1^-/-^* cells over heterozygous cells (Fig. 1h-j). RAD52 siRNA treatment reduced RAD51 foci numbers in irradiated, homozygote S-phase cells, and co-depletion with RADX both restored RAD51 foci numbers and negated the lethality of RAD52 depletion (Fig. 1i & j). Consistent with these data, we found that HR repair outcomes in *Bŗca1^C61G/C61G^ 53bp1^-/-^* cells were reduced by RAD52, RNF168 or BRCA1 siRNA-treatment (Fig. 1k).

We noted that Polθ protein levels were elevated in *53bp1^-/-^* cells compared to WT cells and further increased in *Brca1^C61G/C61G^ 53bp1^-/-^* cells compared to *Brca1^+/+^ 53bp1^-/-^* cells (Fig. 2a, Supplemental Fig. 3a). Upon depletion of Polθ, the survival of cells lacking *53bp1* was reduced to a greater extent with *Bŗca1^C61G/C61G^* than with WT *Brca1* (Supplemental Fig. 3b, Fig. 2b). Sensitivity to Polθ depletion coincided with an increase in RAD51 foci numbers in untreated *Brca1^C61G/ C61G^ 53bp1^-/-^* cells and in IR-treated *Brca1^C61G/C61G^ 53bp1^-/-^* S-phase cells, and also with fewer RAD51 foci in 24 hours after IR (Fig. 2c & d). In DNA repair through TMEJ the 3’ flaps are cleaved, and strands ligated by XRCC1/ligase III^29^. To test whether *Brca1^C61G/C61G^ 53bp1^-/-^ cells* depend on ligase III for survival, we assessed the impact of the ligase I/III inhibitor, L67^30^. We noted no differential impact of L67 on cell survival in otherwise untreated cells (Supplemental Fig. 3c). Further, *Brca1^C61G/C61G^ 53bp1^-/-^* cells showed no increase in TMEJ repair products following repair at a specific DNA break compared to outcomes from *Brca1^+/+^ 53bp1^-/-^* cells (TMEJ products were 19.9% of reads in *Brca1^+/+^ 53bp1^-/-^* and 18.6% of reads in *Brca1^C61G/C61G^ 53bp1^-/-^* cells, Supplemental Fig. 3d). Thus, *Brca1^C61G/C61G^ 53bp1^-/-^* cells are dependent on Polθ to support survival and normal RAD51 kinetics, but do not exhibit dependence on or increased engagement of TMEJ.

**Fig. 2.**
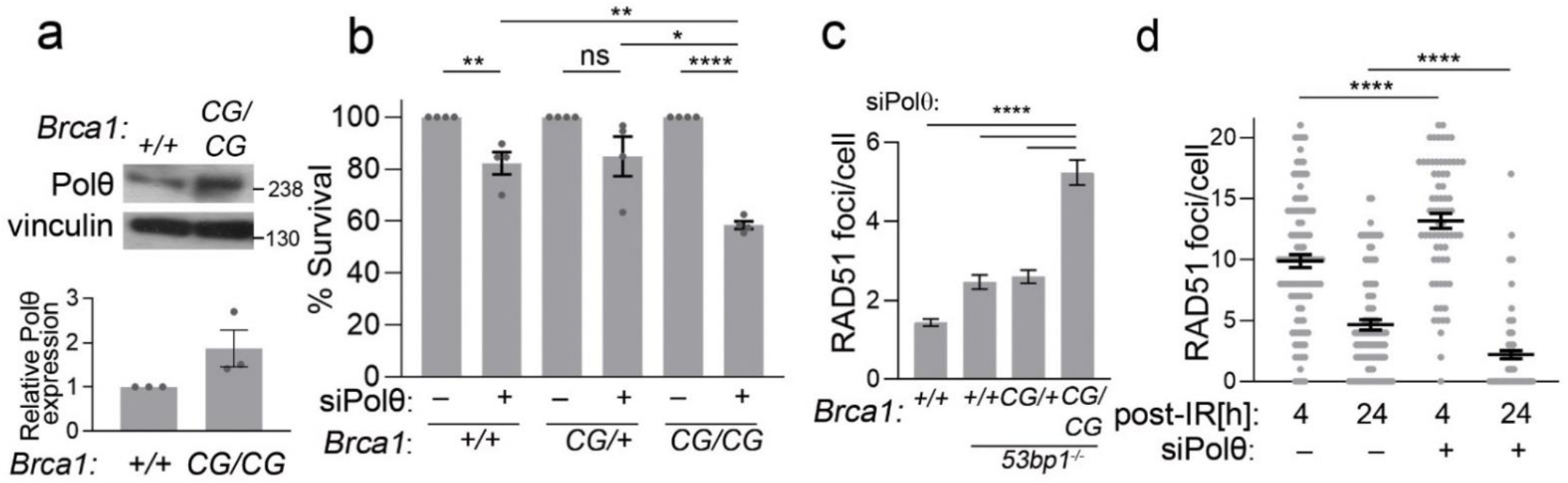
*Brca1^C61G/C61G^ 53bp1^-/-^* cells are sensitive to Polθ depletion. **a**. Western blot of Polθ and quantification of protein levels in the genotypes shown (all are *53bp1^-/-^*). Graph below shows quantification of Polθ relative to loading controls in *Brca1^+/+^ 53bp1^-/-^* and Brca1^C61G/C61G^ 53bp1^-/-^ cells. N=3. Data are mean ± s.e.m. **b**. Colony survival in MEFs of the genotypes shown (all are *53bp1^-/-^*), treated, with NTC siRNA (-) or siRNA targeting Polθ (+), *n*=4 Data are mean ± s.e.m. **c.** Quantification of RAD51 foci, in EdU-positive MEFs of the genotypes shown, which were treated with siRNA to Polθ. *n* = 100 cells from 2 biological replicates, pre condition. Data are mean ± s.e.m. **d**. Quantification of RAD51 foci 4 and 24 hours after 2 Gy IR exposure, in EdU-positive *Brca1^C61G/C61G^ 53bp1^-/-^* cells treated, with NTC siRNA (-) or siRNA targeting Polθ (+). n = 100 cells from 2 biological replicates, per condition. Data are mean ± s.e.m.

We next addressed potential relationships between Polθ, RAD52 and RNF168 in *Brca1^C61G/C61G^ 53bp1^-/-^* cells. Polθ siRNA treatment had negligible impact on RNF168 foci formed at γH2AX-decorated sites (Fig. 3a & b), whereas RAD52 accumulations were increased (Fig. 3c-e). Similarly, treatment with the Polθ inhibitor, ART558, but not treatment with the ligase I/III inhibitor, L67, increased the number of RAD52 foci in *Brca1^C61G/C61G^ 53bp1^-/-^* cells (Supplemental Fig. 4a & b). We titrated RAD52 inhibitor and RNF168 siRNA in irradiated, Polθ depleted *Brca1^C61G/C61G^ 53bp1^-/-^* cells to identify the concentrations able to restore RAD51 foci to control levels (Fig. 3f & g). We then tested the impact of these concentrations on cell survival. Remarkably, low concentrations of RAD52 inhibitor, but not RNF168 siRNA, improved the viability of *Brca1^C61G/C61G^ 53bp1^-/-^* cells treated with Polθ siRNA (Fig. 3h-i). Low concentrations of RAD52 siRNA able to normalise RAD51 foci to control levels following Polθ siRNA treatment similarly restored cell survival (data not shown). *Brca1^C61G/ C61G^ 53bp1^-/-^* cells exhibit decreased replication fork protection after hydroxyurea treatment (Supplemental Fig. 4c), and although the addition of Polθ siRNA or RAD52 inhibition alone did not worsen the defect, their combination improved DNA replication fork stability (Supplemental Fig. 4d). In contrast, the increased replication fork stalling seen in *Brca1^C61G/ C61G^ 53bp1^-/-^* cells was not improved (Supplemental Fig. 4e & f). RAD52 inhibition was also found to suppress chromosome abnormalities induced by Polθ siRNA in metaphases of *Brca1^C61G/C61G^ 53bp1^-/-^* cells (Fig. 3j), and RAD52 siRNA treatment corrected the aberrant RAD51 foci kinetics seen after IR (Supplemental Fig. 4g). RAD52 inhibition similarly improved the survival of WT cells (i.e. *53bp1^+/+^*) depleted for BRCA1 and treated with Polθ inhibitor (Supplemental Fig. 4h). These data suggest RAD52 mediates many of the deleterious features associated with Polθ loss or inhibition in BRCA1-deficient cells.

**Figure. 3.**
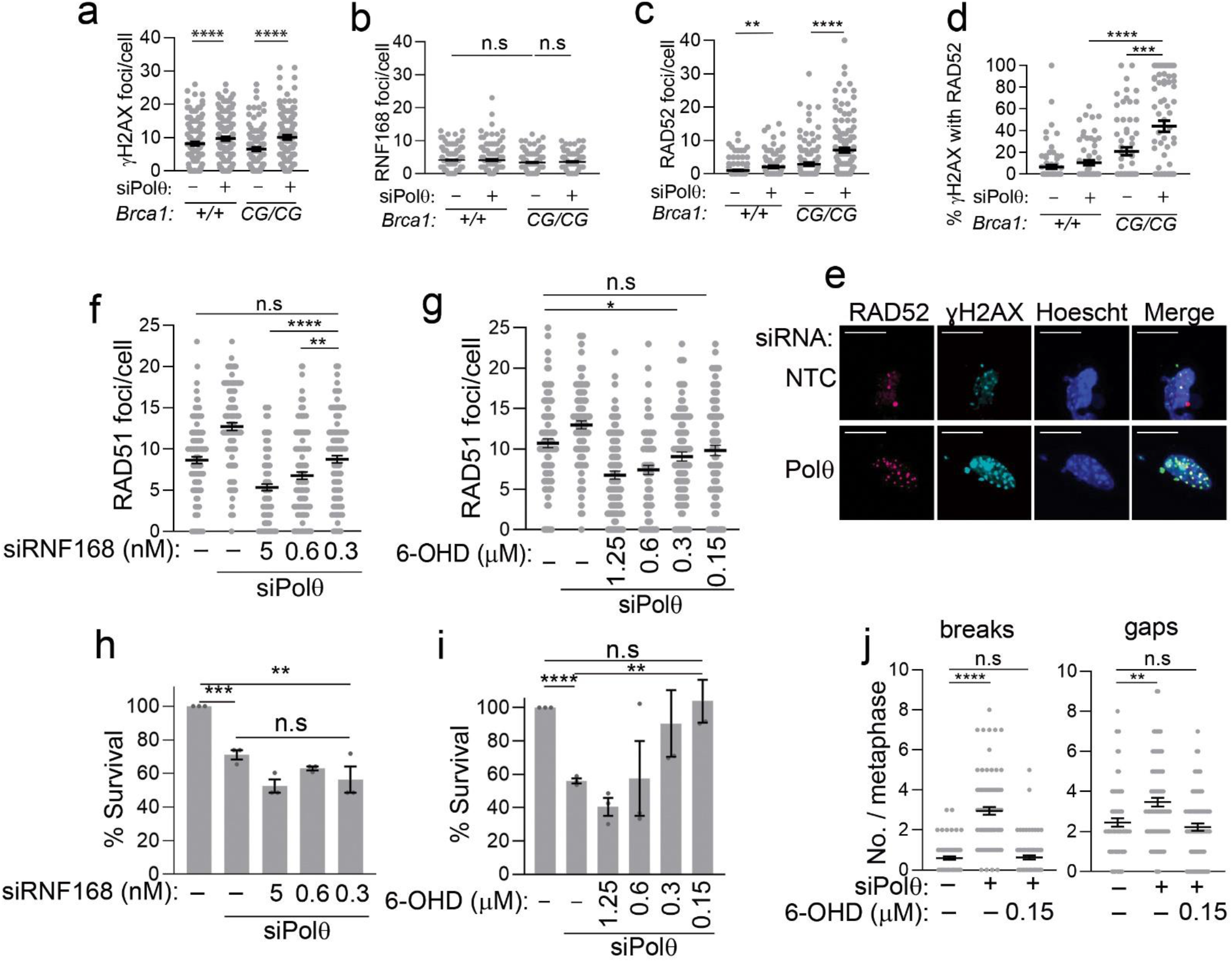
Polθ suppresses RAD52-mediated toxicity. **a to c.** (**a**) γH2AX foci, RNF168 foci (**b**), and flag-RAD52 foci (**c**) in the genotypes shown (all are *53bp1^-/-^*). treated with control (-) or Polθ(+) siRNA. In each experiment *n* = 120 cells from 3 biological replicates, per condition. Data are mean ± s.e.m. **d.** Percentage of γH2AX foci colocalised with RAD52 foci in *Brca1^+/+^ 53bp1^-/-^* cells Vs *Brca1^C61G/C61G^ 53bp1^-/-^* cells treated with control (-) or Polθ(+) siRNA. *n* = 80 cells from 2 biological replicates, per condition. Data are mean ± s.e.m. **e.** Representative images of RAD52 and γH2AX in *Brca1^C61G/C61G^ 53bp1^-/-^* cells treated with control or Polθ siRNA. Scale bars represent 10 μm. **f. and g**. Quantification of RAD51 foci in EdU-positive in *Brca1^C61G/C61G^ 53bp1^-/-^* cells treated with siRNA targeting Polθ and decreasing concentrations of RNF168 siRNA (**f**) and decreasing concentrations of RAD52 inhibitor (**g**) (6-OH-Dopa). refers to treatment with non-targeting control siRNA or vehicle. *n* = 150 cells from 3 biological replicates, per condition Data are mean ± s.e.m. **h. and i**. Colony survival of *Brca1^C61G/C61G^ 53bp1^-/-^* cells treated with targeting Polθ siRNA and the concentrations of RNF168 siRNA (**h**) and RAD52 inhibitor (**i**). refers to treatment with non-targeting control siRNA or vehicle. *n*=3 Data are mean ± s.e.m. **j**. Number of breaks and gaps per metaphase spread in *Brca1^C61G/C61G^ 53bp1^-/-^* cells treated with control (-) or Polθ (+) siRNA in presence or absence RAD52 inhibitor 6-OH-Dopa. n=80-90 metaphases from 3 biological repeats. Data are mean ± s.e.m.

To investigate why Brca*1^C61G/ C61G^ 53bp1^-/-^* cells might be more sensitive to Polθ repression we first investigated C61G-BRCA1:BARD1 interactions. To examine the interaction independently of expression levels we compared human C61G-BRCA1 with the BARD1 disruptive variant, M18T-BRCA1^31^. C61G and M18T substitutions each reduced the ability of BRCA1 to co-purify BARD1, or to form BARD1-induced foci, while the C61G-M18T-BRCA1 double mutant co-purified no detectable BARD1 and had very few BARD1-induced foci (Supplemental Fig. 5), illustrating a reduced, but not lost, interaction between human C61G-BRCA1 and BARD1. Similarly, expression of exogenous murine BARD1(Supplemental Fig. 5d) increased C61G-BRCA1 foci numbers in irradiated *Brca1^C61G/C61G^ 53bp1^-/-^* MEFs (from 6.5 to 12.7/cell) (Fig. 4a & b). This BARD1 expression also increased HR outcomes measured by PCR by 27% (Fig. 4c) and improved cellular resistance to RAD52 and Polθ depletion by 30%, the latter in a BRCA1-dependent manner (Fig. 4d). Expression of BARD1 mutants that prevent BRCA1 (L38R)^31^ or nucleosome (A448T, D700A)^32, 33^ interactions failed to improve C61G-BRCA1 foci or to promote resistance to Polθ depletion (Fig. 4e & f). In contrast, the AAE-BARD1 mutant (F125A, D127A, A128E), which disrupts RAD51 binding^34^, could improve C61G-BRCA1 foci but was less able to promote resistance to Polθ siRNA treatment (Fig. 4e & f). Exogenous WT-BARD1 expression suppressed RAD52 foci formation in cells treated with Polθ inhibitor, ART558, whereas AAE-BARD1 expression did not (Fig. 4g & h). These data suggest BRCA1 recruitment through BARD1 and BARD1:RAD51 interactions contribute to suppression of RAD52 recruitment and cellular resistance to Polθ depletion.

**Fig. 4.**
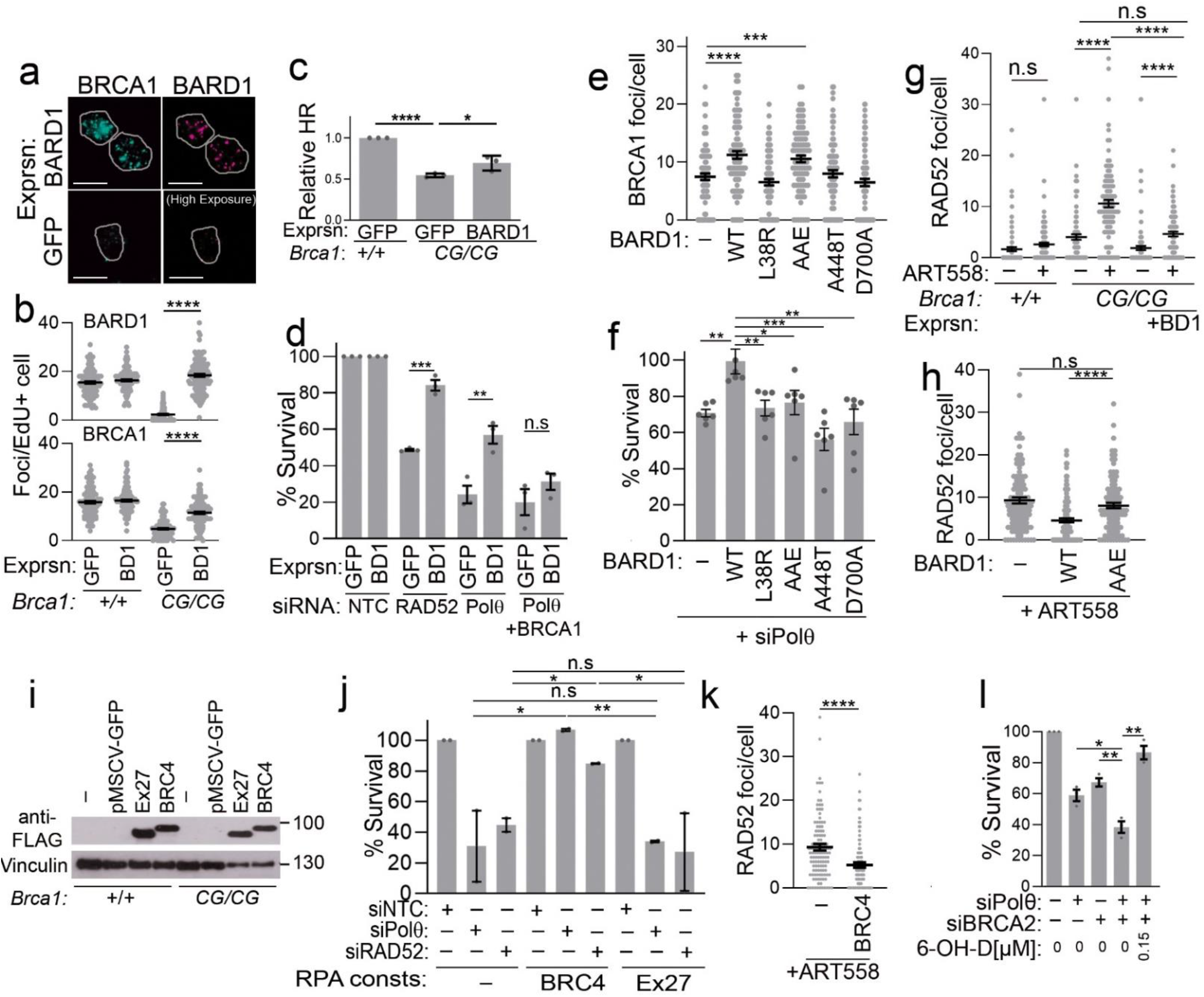
BRCA-RAD51 interactions suppress RAD52 recruitment and Polθ dependency. **a.** Representative images of endogenous C61G-BRCA1 (cyan) in *Brca1^C61G/C61G^ 53bp1^-/-^* cells 2 hours after 2 Gy IR exposure, in EdU-positive cells, with co-expression of murine BARD1 (magenta) or empty vector (GFP). Scale bar is 10 μm. **b**. Quantification of BARD1 (top) and BRCA1 (bottom) foci in the genotypes shown (all are *53bp1^-/-^*). and transfected with empty vector or BARD1 from EdU positive cells following a 2 hour recovery from exposure to 2 Gy IR. n= 110 from 3 biological repeats Data are mean ± s.e.m. **c**. Relative HR-PCR product intensity in the genotypes shown (all are *53bp1^-/-^*). expressing BARD1 or empty vector compared to *Brca1^+/+^ 53bp1^-/-^* cells expressing empty vector. *n*=3 biological replicates (each of 1 technical replicate). Data are mean ± s.e.m. **d.** % Colony survival of *Brca1^C61G/C61G^ 53bp1^-/-^* cells treated with control (NTC), RAD52, Polθ, or Polθ and BRCA1 siRNA and infected with GFP or BARD1 expressing retrovirus. n= 3 biological repeats, data are mean ± s.e.m. **e.** Quantification of BRCA1 foci in *Brca1^C61G/C61G^ 53bp1^-/-^* cells infected with empty, pMSCV-IRES-EGFP (-), WT-BARD1 or BARD1 mutant retrovirus and stained for BRCA1 n= 90, across 3 biological repeats. Data are mean ± s.e.m **f.** % Colony survival in *Brca1^C61G/C61G^ 53bp1^-/-^* cells treated with Polθ siRNA and infected with infected with empty, pMSCV-IRES-EGFP (-), WT-BARD1 or BARD1 mutant retrovirus control n= 6 biological repeats, data are mean ± s.e.m. **g**. Quantification of flag-RAD52 foci in cells of the genotypes shown (all are *53bp1^-/-^*), treated with vehicle or (10 μM) ART558 and infected with empty pMSCV-IRES-EGFP (-) or retrovirus bearing WT-BARD1. n=90 from 3 biological repeats, data are mean ± s.e.m. **h**. Quantification of **F**lag-RAD52 foci in *Brca1^C61G/C61G^ 53bp1^-/-^* cells infected with retrovirus bearing WT-BARD1 or AAE-BARD1 mutant, then treated with 10 μM ART558 for 72h. n= 100 from 3 biological repeats, data are mean ± s.e.m. **i.** Western blot showing expression of RPA-70-BRCA2-Exon27 (Ex27) or RPA-70-BRCA2-BRC4 (BRC4) in cells of the genotypes shown (all are *53bp1^-/-^*). **j.** % Colony survival in *Brca1^C61G/C61G^ 53bp1^-/-^* cells treated with non-targeting control (NTC), Polθ and RAD52 siRNA and infected with empty pMSCV-IRES-EGFP retrovirus or those expressing RPA-70-BRCA2-BRC4 (BRC4) or RPA-70-BRCA2-Exon27 (Ex27). n= 2, 2 biological repeats. Data are mean ± s.e.m. **k**. Quantification of flag-RAD52 foci in *Brca1^C61G/C61G^ 53bp1^-/-^* cells infected with retrovirus bearing empty vector or RPA-70-BRCA2-BRC4 (BRC4), then treated with 10 μM ART558 for 72h. n= 90 from 3 biological repeats, data are mean ± s.e.m. **l**. % Colony survival of *Brca1^C61G/C61G^ 53bp1^-/-^* cells treated with control (-) or siRNA targeting Polθ, BRCA2 or both in presence of RAD52 inhibitor 6-OH-Dopa or vehicle. n= 3, 3 biological repeats Data are mean ± s.e.m.

A function of BRCA1 is to promote the accumulation of PALB2-BRCA2 to sites of damage, and BRCA1-BARD1 expression is insufficient to prevent sensitivity to Polθ loss in *BRCA2* deficient cells^2, 3^. To explore whether BRCA2: RAD51 interactions might contribute to resistance we expressed one of the BRCA2 BRC repeats, BRC4, and Exon27 regions of BRCA2 fused to the RPA subunit RPA-70 (Fig. 4i). BRCA2-BRC4 aids the exchange of RAD51 with RPA^35^ and the C-terminus contacts oligomerised RAD51 to support its stability^36, 37^. Expression of RPA-BRC4, but not RPA-Exon27 overcame the toxicity of Polθ and RAD52 siRNAs in *Brca1^C61G/C61G^ 53bp1^-/-^* cells (Fig. 4j) and RPA-BRC4 expression could suppress the formation of RAD52 foci on Polθ inhibitor treatment (Fig. 4k). These data suggested that the sensitivity of BRCA2 deficient cells to Polθ loss might also be mediated by RAD52. We depleted BRCA2 in *Brca1^C61G/C61G^ 53bp1^-/-^* cells and observed a further increase in their susceptibility to Polθ siRNA treatment. Remarkably this was suppressed by the addition of RAD52 inhibitor (Fig. 4l), suggesting that RAD52 is required for toxicity of targeting Polθ in cells with reduced BRCA2.

*POLQ* knock out and *53bp1* loss is synthetic lethal^1, 4^, although *53bp1^-/-^* cells are less sensitive to Polθ siRNA or Polθ inhibitor than *Brca1^C61G/C61G^ 53bp1^-/-^* cells (Fig. 2b, Supplemental Fig. 6a). Since we observed increased RAD52 foci in *53bp1^-/-^* cells following Polθ depletion, and restoration of *Brca1^C61G/C61G^ 53bp1^-/-^* cell viability on RAD52 inhibition (Fig. 3c & i), we predicted that the susceptibility of *53bp1^-/-^* cells to Polθ inhibition might also be mediated by RAD52. Indeed, we saw that co-treatment with low concentrations of RAD52 inhibitors^28, 38^ suppressed the toxicity of Polθ inhibition in *53bp1^-/-^* cells (Supplemental Fig. 6b and c).

In these studies, we find that *Brca1^C61G/C61G^ 53bp1^-/-^* cells exhibit several features of HR competency and require the mutant BRCA1, and wild-type RNF168 and RAD52 to support RAD51 accumulations, HR, and cell viability. Heterozygote *Brca1^C61G/+^ 53bp1^-/-^* cells do not depend on RAD52. RAD52-dependency has been noted in HR defective cells entirely lacking *BRCA1, PALB2* or *BRCA2* ^27^, so we propose that BRCA1 dysfunction draws first on RNF168, then on RAD52 to maintain HR and cell survival.

The molecular foundations of synthetic lethality between HR gene loss or *53BP1* loss and targeting Polθ have not been clear. Here we find that RAD52 mediates much of the toxicity including promoting the occurrence of chromosome breaks, aberrant RAD51 kinetics and increased cell death. RAD52 accumulations at sites of DNA damage are increased in mutant cells following Polθ suppression, and we find that BRCA1-BARD1 and BRCA2 protein contacts with RAD51 are implicated in suppressing RAD52 accumulations and repressing cellular sensitivity to Polθ inhibition. These findings are consistent with the ability of BRCA2 to suppress single-strand annealing^15^, which is mediated by RAD52. They suggest that RAD51:ssDNA nucleoprotein formation or stabilization suppresses RAD52 accumulations in the absence of Polθ.

We find that cells with both *Brca1* mutation and *53bp1* loss are more sensitive to Polθ suppression than *53bp1^-/-^* cells, consistent with recent reports^7, 8^. Nevertheless, the sensitivity of *53bp1^-/-^* cells also requires RAD52. Polθ helicase activity competes with RPA^6^ and cells treated with a Polθ inhibitor exhibit greater RPA phosphorylation and extended resection^7^. Since extended resection and increased RPA promote RAD52 recruitment^39, 40^ this may explain how Polθ suppresses RAD52. We speculate that excessive RAD52, whether in HR-deficient or *53bp1* deficient cells, in turn leads to inappropriate recombination. We do not discount the possibility that other mechanisms contribute to lethality in Polθ inhibited cells. CRISPR-Cas9 screens have revealed that knockout of many DNA repair genes result in sensitivity to POLQ loss, including deletion of RAD52 itself ^4^, so the relationship described herein is not universal. Our finding of a hierarchy of support mechanisms for HR, uncovered by the investigation of the C61G-BRCA1 hypomorph, and the requirement for RAD52 in both HR and *53bp1* deficient cells is consistent with the spectrum of responses to Polθ inhibitor seen in BRCA1 and BRCA2-deficient cancer models recently described^7, 8^.

**Materials and Methods** are in the Supplementary Information file.

## Author Contributions

KS (Colony Assays, Immunofluorescence, RAD51 foci analysis, metaphase spreads, DNA fibre assay) GR (Mass Spec preparations, immunofluorescence, HR measurements, construct design RAD52, hBRCA1, colony assays, Western blots), EA (Colony assays, Immunofluorescence, Western blots). LC (murine BARD1 constructs), AJG (mouse colony). ME (antibody development, MEF generation). GS (ART558 generation), AB & CW, (sequencing and bioinformatic analysis). JB: Genotyping. JRM (supervision and project direction). All authors read and reviewed the manuscript.

## Acknowledgements

ART558 was a kind gift of Graeme Smith Artios Pharma Ltd. Cambridge, U.K. SV40 1: pBSSVD2005 was a gift from David Ron (Addgene plasmid # 21826; http://n2t.net/addgene:21826; RRID:Addgene_21826). Anti murine BARD1 antibody was a gift from Richard Baer (Columbia). 53BP1 animals were a gift from Andre Nussenzweig, NIH (National Institutes of Health), Bethesda. GR, JB and KS are funded by Cancer research, UK (C8820/A28283), ME and KS by Breast Cancer Now, UK (2015MayPR499), EA by the Midlands Integrative Biosciences Training Partnership, funded by the Biotechnology and Biological Sciences Research Council, UK. AJG by the Wellcome Trust (206343/Z/17/Z), A.B and C.W by Cancer Research UK (C31641/A23923). We thank the microscopy and Imaging services (MISBU) at the University of Birmingham Technology Hub for their assistance and maintenance of equipment. We sincerely apologise to colleagues whose excellent and relevant work was not referenced due to journal space restrictions.

## Supplemental Figures and Legends

**Supplemental Fig. 1.**
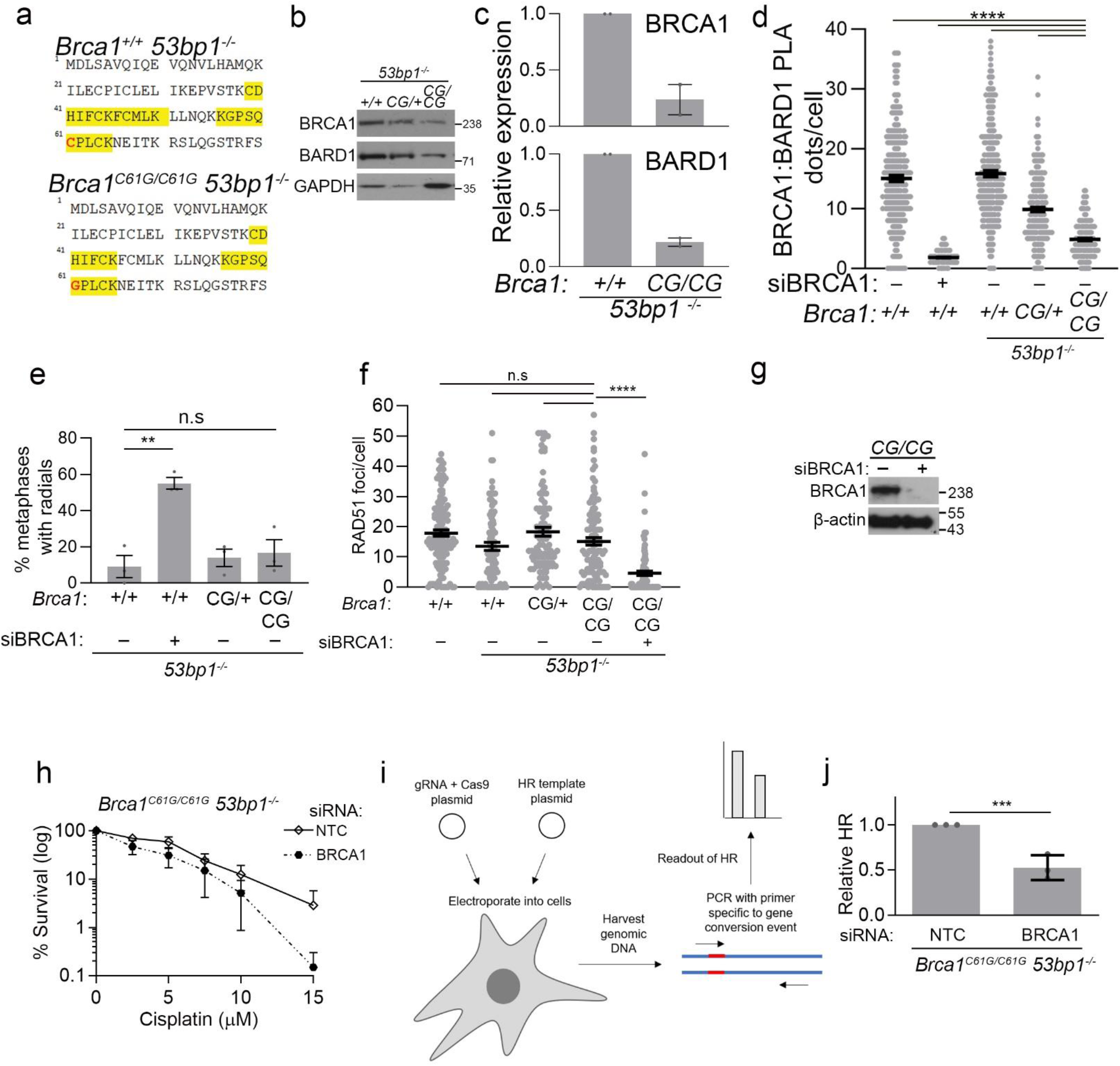
*Brca1^C61G/C61G^ 53bp1^-/-^* cells support HR. **a**. BRCA1 peptides identified by mass spectrometry from *Brca1^+/+^ 53bp1^-/-^ and Brca1^C61G/C61G^ 53bp1^-/-^* mouse embryonic fibroblasts. **b**. Representative western blot analysis of BRCA1, BARD1 and 53bp1 protein levels in *MEFs*. Expression of GAPDH was used as a loading control. **c**. BRCA1–BARD1 protein levels in *Brca1^+/+^ 53bp1^-/-^ and Brca1^C61G/C61G^ 53bp1^-/-^* cells, relative to loading controls and wild-type protein. *n* = 2, data are mean ± s.e.m. **d**. Quantification of proximity linked ligation assay foci (PLA) between BRCA1 and BARD1 in cells treated with 2 Gy irradiation and with NTC siRNA (-) or siRNA targeting BRCA1 (+). *n* = 200-300 cells from 5 biological replicates; bars depict median ± s.e.m. A two-sided unpaired *t*-test was used to calculate all *P* values. **e**. Percentage of metaphases that show one or more radial chromosomes in MEFs of the genotypes shown, treated with NTC siRNA (-) or siRNA targeting BRCA1(+). n=3 data are mean ± s.e.m. **f.** Quantification of RAD51 foci following 16 hours 5 μM cisplatin exposure in EdU-positive MEFs of the genotypes shown, treated with NTC siRNA (-) or siRNA targeting BRCA1(+). *n* = at least 100 cells from 3 biological replicates, respectively. Data are mean ± s.e.m. **g.** Representative western blot showing knockdown of BRCA1 in *Brca1^C61G/ C61G^ 53bp1^-/-^* cells. **h**. Colony survival following 16 h treatment with cisplatin, measured in cells in *Brca1^C61G/C61G^ 53bp1^-/-^* cells treated with non-targeting control siRNA (NTC) or BRCA1 siRNA transfection, *n*=3, data are mean ± s.e.m. i. Illustration of the PCR-based measure for HR outcomes. After electroporation with template DNA and plasmid containing CAS9 and Rosa26 guide RNA the relative PCR product intensity of HR-specific *Vs* locus specific PCR product is assessed. **j**. Quantification of HR-PCR product between *Brca1^C61G/C61G^ 53bp1^-/-^* and *Brca1^+/+^ 53bp1^-/-^* cells. n=3 biological replicates data are mean ± s.e.m.

**Supplemental Fig. 2.**
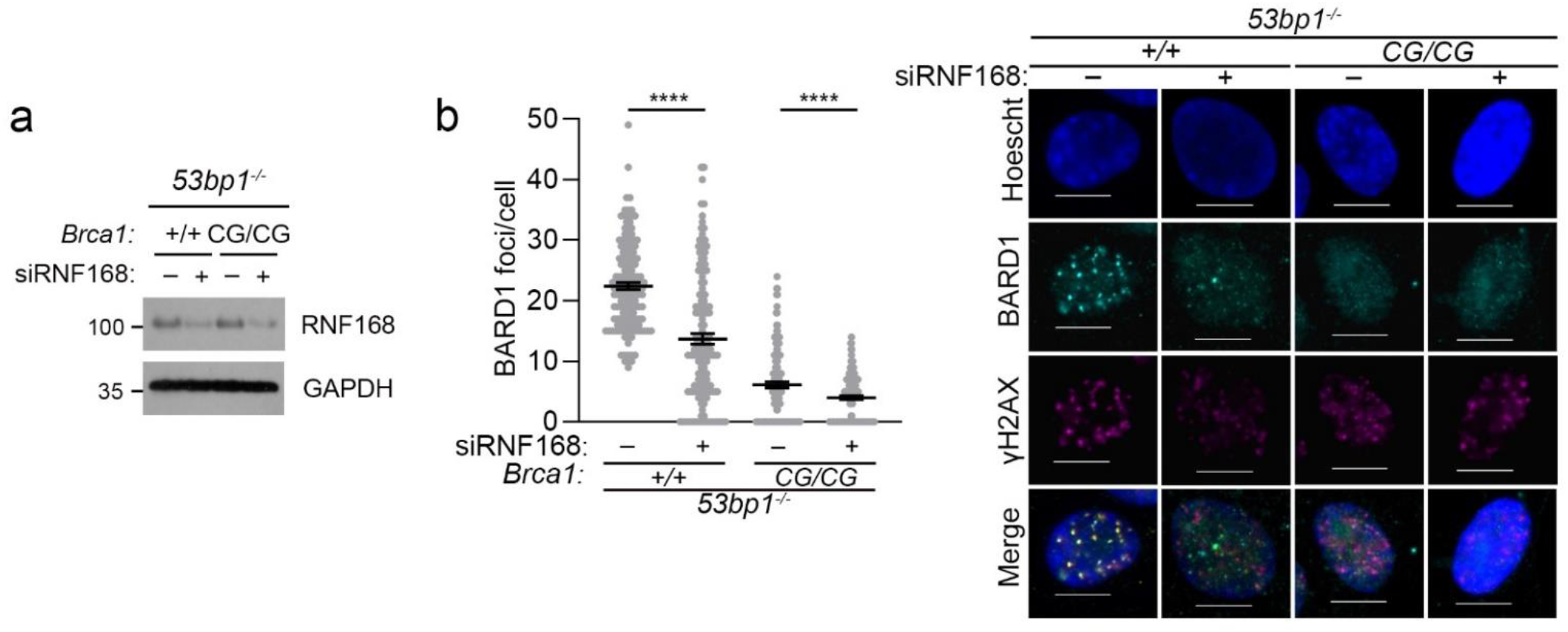
Impact of RNF168 depletion on BARD1 localisation. **a.** Western blot analysis of RNF168 protein levels following control (-) or RNF168(+) targeting siRNA treatment. **b.** Quantification of BARD1 foci 3 hours after 2 Gy IR exposure, in EdU-positive MEFs of the genotypes shown and which were treated with NTC siRNA or RNF168 siRNA. *n* = 150 cells across 3 biological replicates. Data are mean ± s.e.m. Images right show representative images of BARD1 foci in the genotypes shown with and without RNF168 depletion. Scale bar is 10 μm.

**Supplemental Fig. 3.**
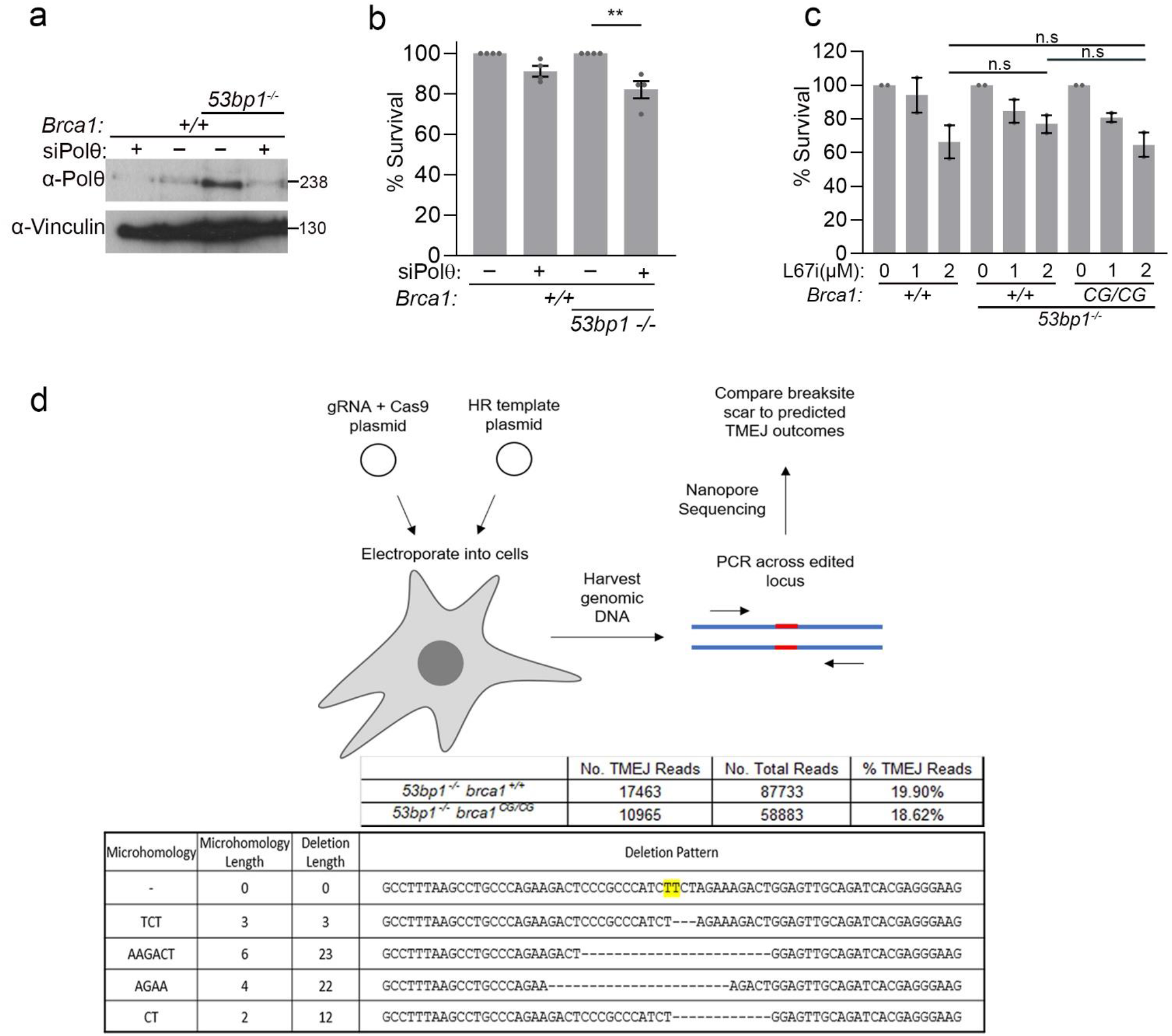
*Brca1^C61G/C61G^ 53bp1^-/-^* cells are sensitive to Polθ depletion. **a**. Western blot of Polθ in genotypes shown treated with control (-) siRNA or siRNA to Polθ (+). **b.** Colony survival in MEFs of the genotypes shown treated with control (-) or Polθ (+) siRNA n= 4, 4 biological repeats Data are mean ± s.e.m. **c**. Colony survival in MEFs of the genotypes shown treated with L67 inhibitor n= 2,2 biological repeats Data are mean ± s.e.m. **d.** Illustration of TMEJ readout. The upper table shows the % of reads containing predicted TMEJ outcomes from the total number of reads. The lower table shows the top five predicted TMEJ outcomes. DSB is induced between the two highlighted bases.

**Supplemental Fig. 4.**
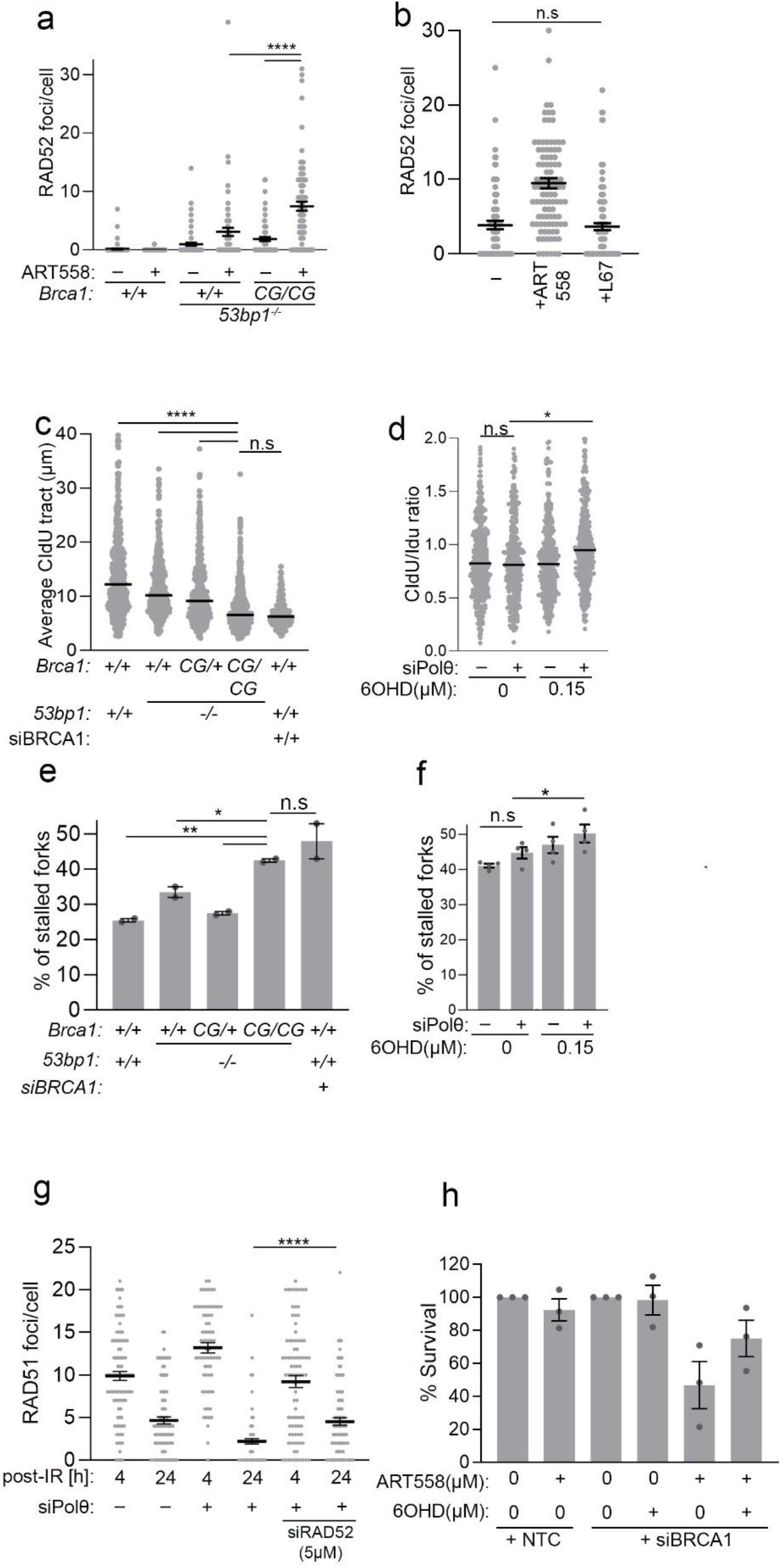
Polθ counters RAD52. **a.** Quantification of flag-RAD52 foci in cells of the genotypes shown and treated with vehicle or 10 μM Polθ inhibitor ART558 for 72 hours. n = 70 cells from 2 biological replicates. Data are mean ± s.e.m. **b**. Quantification of flag-RAD52 foci in *Brca1^C61G/C61G^ 53bp1^-/-^* cells treated with vehicle, 10 μM Polθ inhibitor ART558 or 2 μM ligase I/III inhibitor L67 for 72 hours. n = 90 cells from 3 biological replicates. Data are mean ± s.e.m. **c**. Average CIdU tract length from cells with the illustrated genotypes treated with hydroxyurea (5 mM, 3 h). N = at least 200 fibre from 3 replicates. Bars = median **d**. IdU:CldU ratios from *Brca1^C61G/C61G^ 53bp1^-/-^* cells treated with Polθ siRNA, and or RAD52 inhibitor, 6-OH-Dopa and hydroxyurea (5 mM, 3 h). n=at least 350 fibres from 3 replicates. Bars=median **e**. % cells with first label terminations from cells with the illustrated genotypes. n=at least 800 fibre from 2 replicates. Data are mean ± s.e.m. **f**. % cells with first label terminations from *Brca1^C61G/C61G^ 53bp1^-/-^* cells treated with Polθ siRNA, and or RAD52 inhibitor, 6-OH-Dopa. n = at least 2500 fibre from 4 replicates. Data are mean ± s.e.m. **g.** Quantification of RAD51 foci 4 and 24 hours post 2 Gy IR exposure, in EdU-positive in *Brca1^C61G/C61G^ 53bp1^-/-^* cells treated with siRNA targeting Polθ (+), or control siRNA (-) with or without siRNA targeting RAD52. n = 100 cells from 2 biological replicates, respectively. Data are mean ± s.e.m. **h.** % Colony survival in wild-type cells treated with non-targeting control (NTC) or BRCA1 siRNA, with or without ART558 (10 μM) or 6OHD (0.15 μM) for 72h n = 3, 3 biological repeats. Data are mean ± s.e.m.

**Supplemental Fig. 5.**
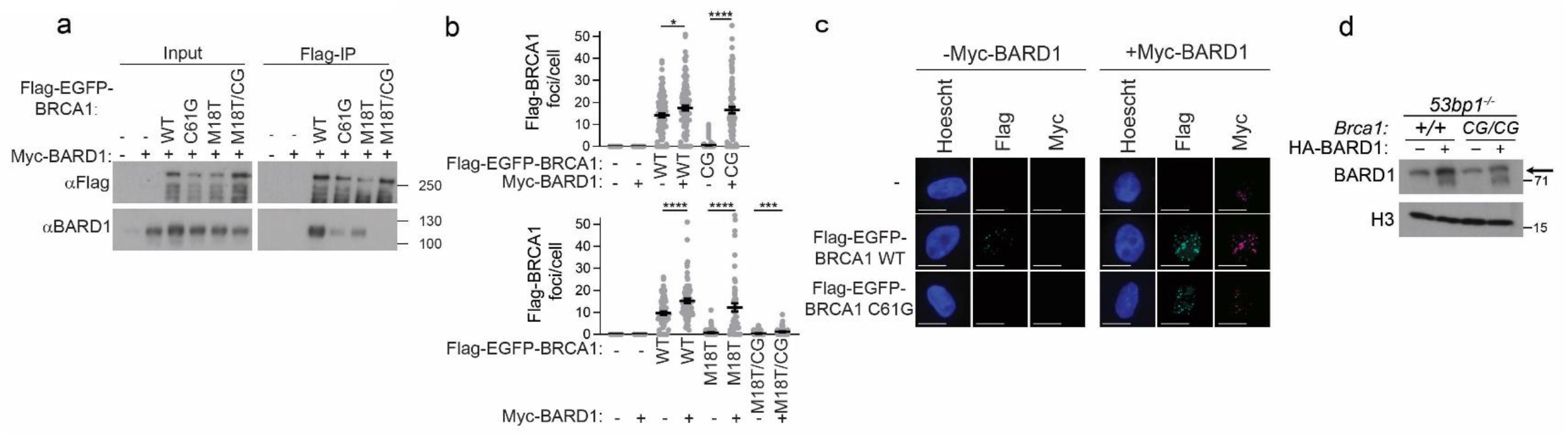
BARD1 overcomes sensitivity of *Brca1^C61G/C61G^ 53bp1^-/-^* cells to Polθ depletion. **a.** Representative blot of Flag immunoprecipitation of human Flag–eGFP-tagged BRCA1 variants from U20S cells expressing, or not exogenous human BARD1. **b.** Quantification of Flag–eGFP-tagged BRCA1 variant foci in cells with and without co-expression of human Myc-tagged BARD1 following a 2-hour recovery from 2 Gy IR. n = 120 from 3 biological repeats (upper panel) and n = 60 from 2 biological repeats (lower panel). Data are mean ± s.e.m **c.** Representative images of Flag–eGFP-tagged BRCA1 foci and Myc-tagged BARD1 foci from U2OS cells following a 2-hour recovery from 2 Gy IR. Scale bars represent 10 μm. **d**. Expression levels of BARD1 in the MEF genotypes shown with (+) and without (-) co-expression of murine BARD1.

**Supplemental Fig. 6.**
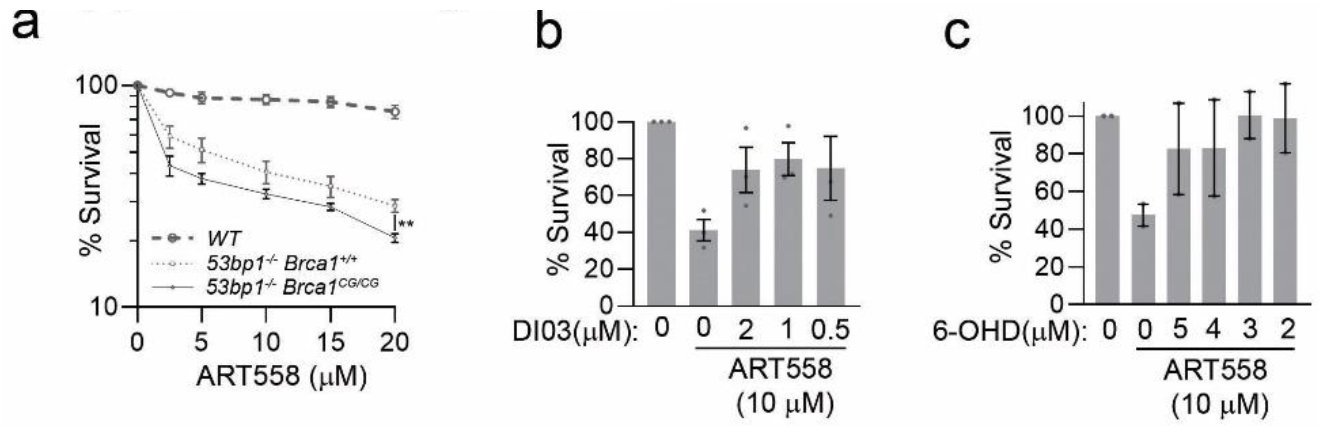
Synthetic lethality between 53BP1 loss and Polθ inhibition is suppressed by RAD52 inhibition. **a.** % Colony survival in wild-type (WT), *53bp1^-/-^* (*Brca1^+/+^*) and *Brca1^C61G/C61G^ 53bp1^-/-^* cells treated with increasing doses of ART558 μM for 72h n = 6, 6 biological repeats Data are mean ± s.e.m. **b**. % Colony survival in *53bp1^-/-^* cells treated with 10 μM ART558 and the decreasing dose of RAD52 inhibitor DI03. n= 3, 3 biological repeats Data are mean ± s.e.m. **c.** % Colony survival in *53bp1^-/-^* cells treated with 10 μM ART558 and the decreasing dose of RAD52 inhibitor 6-OH-Dopa. n = 2, 2 biological repeats Data are mean ± s.e.m.

