## Supplemental Materials and Methods for "RAD52 underlies the synthetic-lethal relationship between BRCA1/2 and 53BP1 deficiencies and DNA polymerase theta loss"

**Animal statement.** The generation of *Brca1* C61G allele is previously described[1]. Mice with the Trp53bp1+/- [2] allele were obtained from the NIH (Bethesda). The Research Ethics Committee for animal experimentation at the University of Birmingham, UK reviewed and the Home Office approved all the work included in this manuscript. All *in vivo* experiments were performed under the UK Animals (Scientific Procedures) Act 1986 Home Office regulations under the authority of PPL70 /8013.

***Cell line maintenance and Generation of Mouse Embryonic Fibroblasts***. $Brca1^{C61G/+}$ $53Bp1^{-/-}$ male and female animals were mated to generate littermates of required *Brca1* genotypes. Pregnant mice were euthanised 13.5 days after mating and the embryos were dissected into media to allow fibroblasts to grow out. MEFs were immortalised by transduction with the SV40 large T antigen (pBsSVD2005, AdGene) using FuGENE (Promega). MEFs, HEK293 and U20S cells were maintained Dulbecco's modified Eagles medium (DMEM) supplemented with 10 % fetal bovine serum (FBS) and 1 % penicillin and streptomycin.

***Mass spectrometric analysis.*** Mouse BRCA1 was immunoprecipitated with the mBRCA1-N-terminal antibody (C40) and the precipitate was briefly run into a polyacrylamide Tris Acetate gradient gel (Invitrogen). The sample was excised into 10 slices, which were washed with 25 mM ammonium bicarbonate followed by acetonitrile. Following this, samples were reduced with 10 mM dithiothreitol at 60 °C followed by alkylation with 50 mM iodoacetamide at RT. Subsequently, samples were digested with trypsin (Promega) at 37 °C for 4 h. Finally, samples were quenched with formic acid and the supernatant was analysed directly without further processing.

Each gel digest was analysed by nano LC/MS/MS with a Waters NanoAcquity HPLC system interfaced to a ThermoFisher Q Exactive. Peptides were loaded on a trapping column and eluted over a 75 µm analytical column at 350 nL/min; both columns were packed with Luna C18 resin (Phenomenex). The mass spectrometer was operated in data-dependent mode, with MS and MS/MS performed in the Orbitrap at 70,000 FWHM and 17,500 FWHM resolution, respectively. The fifteen most abundant ions were selected for MS/MS. Data were searched using a local copy of Mascot searching against a Swissprot Mouse database (forward and reverse appended with common contaminants and BRCA1-C61G). The peptide tolerance was set to 10 p.p.m. and the fragment ion tolerance was set to 0.02 Da. A maximum number of two missed cleavages by trypsin were allowed and carbamidomethylated cysteine and oxidised methionine were set as fixed and variable modifications, respectively.

***Western blotting.*** For a full list of antibodies, see Supplementary Table 1**.** Samples were run on SDS–PAGE protein gels and transferred to an Immobilon-P PVDF membrane. Following the transfer, membranes were blocked in 5 % Marvel milk in PBS containing 0.1% Tween (PBStw), or in 5% BSA with PBStw, for 1 h before incubation with primary antibody at 4 °C for 16 h. Blots were washed in PBStw and then transferred into secondary horseradish peroxidase (HRP)-conjugated antibodies in 5 % Marvel milk for 1 h. Blots were washed in PBStw before probing with 1:1 EZ-ECL mix (Biological Industries). Blots were exposed to X-ray film (WolfLabs) and developed using the Xograph Compact X4 developer. Densitometry calculations were performed using ImageJ[3].

***Immunofluorescence staining.*** Cells were plated at a density of $5 \times 10^4$ cells $mL^{-1}$ in 24-well plates on circular glass coverslips (13-mm diameter). Cells were then treated as described. For cells subject to EdU-staining, these were incubated with EdU at a final concentration of 10 µM for 10 min before fixing and staining was carried out as detailed in the Click-iT EdU Imaging Kits (Life Technologies). Cells were pre-extracted by incubation with ice- cold 0.5 % Triton X-100 in PBS on ice for 5 min before fixation with 4 % PFA. Once fixed the cells were permeabilised for a further 30 min using 0.5 % Triton X-100 in PBS before incubation with blocking solution (10% FCS in PBS for 30 min). Cells were then incubated with primary antibodies in 10% FCS in PBS at 4C overnight. The following day, cells were washed 3x with PBST before incubation with Fluorescent secondary antibody (1:2,000) for 2 h. Cells were then washed three times in PBST and the DNA was stained using Hoechst at a 1:20,000 concentration for 5 min. Excess Hoechst was removed by washing with PBS and coverslips were mounted onto Snowcoat slides using Immunomount mounting medium. For a full list of antibodies, see Supplementary Table 1. Immunofluorescence staining was imaged using a Leica DM6000B microscope with a HBO lamp with a 100-W mercury short-arc UV bulb and four filter cubes, A4, L5, N3 and Y5, which produce excitations at wavelengths 360, 488, 555 and 647 nm, respectively.

***Proximity linked ligation Assay.***MEFs were seeded at $4 \times 10^4$ cells ml−1 onto poly-l-lysine-coated coverslips and irradiated with 2 Gy, before recovery for 3 hour. Cells were pre-extracted for 5 min on ice with pre-extraction buffer (20 mM NaCl, 3 mM $MgCl_2$, 300 mM sucrose, 10 mM PIPES, 0.5% Triton X-100) and fixed in 4% PFA for 10 min before blocking in 5% BSA for 16 h. Blocking medium was removed cells were then incubated with the primary antibodies in 5% FCS in PBS for 1 h at room temperature. After incubation with primary antibodies, cells were washed 2 x 5 min in wash buffer A (Sigma) and subsequently incubated with the MINUS or PLUS PLA probes (Sigma Duolink PLA kit) for 1 h at 37 °C in a warm foil-covered box. Cells were then washed twice for 5 min with wash buffer A (Sigma Duolink PLA kit) and incubated with the Sigma Duolink ligation kit (1× ligation buffer, ligase enzyme) for 30 min at 37 °C. Cells were washed twice for 5 min with wash buffer A and incubated for 100 min at 37 °C with the Sigma Duolink amplification kit (1× amplification buffer, polymerase enzyme). subsequently, the cells were washed for 10 min with wash buffer B (Sigma Duolink PLA kit) at room temperature, incubated 5 min with Hoechst and washed again with wash buffer B twice for 5 minutes. Finally, cells were washed for 1 min with 0.01 % wash buffer B and coverslips were mounted onto Snowcoat slides using Immunomount mounting medium.

**Colony survival assays.** MEFs were plated in 24-well plates at $4 \times 10^4$ cells per ml and treated according to the experiment performed. Cells were trypsinised in 100 µl 1× trypsin and resuspended in 900 µl media. Cells were plated on 6-well plates at limiting dilutions and incubated for a further 7 days at 37 °C at 5 % CO2. Once colonies had grown they were stained with 1 % methylene blue in 50 % ethanol and counted. For a full list of DNA-damaging agents and inhibitors, see Supplementary Table 4.

**Generation of stable cell lines.** Stable cell lines were generated from Flp-In U2OS cells that were co-transfected with human BRCA1 cDNA variant in the pcDNA5/FRT/TO vector, and with the Flp recombinase cDNA in the pOG44 vector. Control transfections were carried out without the pOG44 recombinase. Two days after transfection, cells were selected with 100 µg mL−1 hygromycin, the culture medium was replaced every 2 to 3 days and cells were selected for approximately 2 weeks. After selection, cells were treated with 2 µg ml−1 doxycycline for 72 h to induce expression of exogenous Flag-eGFP-BRCA1.

**Plasmid and siRNA transfection.** FuGENE 6 (Roche) was used as a reagent to transfect cells with DNA plasmids. The ratio used was 4:1 FuGENE (µl):DNA (µg), following the manufacturer's guidelines. siRNA transfections were carried out using the transfection reagent Dharmafect1 (Dharmacon) following the manufacturer's instructions. For a full list of siRNA sequences see Supplementary Table 2.

**Retrovirus production and infection** HEK293T Platinum E cells were transfected with pMSCV-IRES-GFP containing different BARD1 variants or RPA fusions using FuGENE-6 following manufacturer's instructions. Culture media was collected 60 h later and filtered through a 0.45 mm filter. MEFs were infected with retroviral containing media supplemented with 4 µg/mL polybrene (Sigma).
The RPA-70 constructs were generated after previously described BRCA2 fusion approaches[4], $BRCA2_{Exon\ 27}$ corresponds to ALDFLSRLPLPPPVSPICTFVSPAAQKAFQPPRSCG (human BRCA2 residues 3,270-3,305). NLS-Ex27-RPA-flag-stop BRC4 corresponds to EKIKEPTLLGFHTASGKKVKIAKESLDKVKNLFDE (human BRCA2 residues 1,514-1,548). NLS-BRC4-RPA70-Flag-STOP

**Fibre labelling and spreading.** Cells were seeded in 6 $cm^2$ plates and treated with thymidine analogues CldU and IdU. To monitor stability of nascent DNA, cells were incubated at 37 °C with CldU for 20 mins at a final concentration of 25 µM, followed by incubation with IdU (250 µM) for 20 mins and then with 5 mM HU for 3 hours. To monitor CldU fibre lengths, cells were incubated at 37 °C with CldU for 20 mins at a final concentration of 25 µM and then with 5 mM HU for 3 hours. To monitor replication fork restart, cells were incubated at 37 °C with CldU for 20 mins at a final concentration of 25 µM and then with 1 mM HU for 1 hour. The HU was then washed out with 3x PBS washes and cells were incubated for a further 40 mins in media containing 250 µM IdU at 37°C.

After incubation with thymidine analogues, cells were washed 2x with ice-cold PBS for 5 minutes with rotation then trypsinised, resuspended in 1 mL of PBS and counted. The optimal concentration is $50 \times 10^4$ cells/ml and thus cells were adjusted to such concentration. 2 µl of the cell sample was placed on Snowcoat microscope slides and allowed to slightly dry for 7 mins. Then 7 µl of spreading buffer (200 mM Tris pH7.4, 50 mM EDTA, 0.5 % SDS) was mixed with the sample and incubated for 2 mins to lyse the cells. In order to spread the sample down the slide, slides were gradually tilted and once the sample had reached the bottom of the slide, they were allowed to dry for 2 mins. Finally, slides were fixed in a 3:1 ratio of Methanol: Acetic acid for 10 mins before leaving slides to air dry for 5-10 mins. Dried slides were stored at 4°C till staining.

***Fibre Immunostaining.*** After fibre spreading slides were washed twice for 5 minutes with 1 mL $H_2O$ and rinsed with 2.5 M HCl before denaturing DNA with 2.5 M HCl for 1 hour 15 mins. Slides were then rinsed 2 x with PBS and washed for 5 minutes in blocking solution (PBS, 1 % BSA, 0.1 % Tween20). Slides were incubated for 1 hour in blocking solution. After blocking, each slide was incubated with 115 µl of primary antibodies, Rat αBrdU (AbD Serotec/Abcam) to detect CldU used at a concentration of 1:2000 and Mouse αBrdU (Becton Dickinson) to detect IdU used at 1:500. Slides were covered with large coverslips and incubated with the antibodies for 1 hour. After incubation with the primary antibody, slides were rinsed 3x with PBS and then incubated for 1 min, 5 mins and 30 mins, with blocking solution. After rinsing and washing, slides were incubated with 115 µl of secondary antibodies (α-Rat AlexaFluor 555 and α-Mouse AlexaFluor 488) in blocking solution, at a concentration of 1:500, covered with a large coverslip for 2 hours. Slides were rinsed 3x with PBS and incubated with blocking solution for 1 min, 5 mins and 30 mins. After again rinsing 2 x with PBS, immunomount mounting media was added to the slide and a large coverslip placed over and left to dry. Coverslips were then stored at –20 °C for microscopy analysis. It is important to point out that during this process slides must be kept protected from light.

**Metaphase spreads.** MEFs were treated with 5 mM HU for 3 h and then incubated with colcemid (0.01 µg mL−1) for 16 h. Cells were trypsinised and centrifuged at 1,200 r.p.m. for 5 min. The supernatant was discarded and cells were resuspended in PBS and centrifugated again. Five millilitres of ice-cold 0.56 % KCl solution was added, and cells were incubated at 37 $^{0}$C for 15 min before centrifuging at 1,200 r.p.m. for 5 min. The supernatant was discarded and the cell pellet was broken before fixation in 5 ml of ice-cold methanol: glacial acetic acid (3:1). Excess of fixation agents were removed and 10 µl of the cell suspension was dropped onto an acetic-acid-humidified slide. Slides were allowed to dry for at least 24 h and then stained with Giemsa solution (Sigma) diluted 1:20 for 20 min. Slide mounting was performed with Eukitt (Sigma).

**CRISPR/Cas9 HR assay.** Adapted from[5]. MEFs (2 x $10^{6}$/condition) were electroporated using the 100 µL Neon electroporation system (1350 V, 30 ms, 1 pulse) to introduce 10 µg of pX459 V2.0 containing Cas9 and a gRNA targeting Rosa26 locus, alongside 10 µg of pUC57 containing a Rosa26 HR template with a 4 bp edited sequence. Following electroporation, cells were plated into antibiotic free media and allowed to recover. Cells were harvested 72 h later, and genomic DNA isolated using a DNEasy Blood and Tissue kit, following manufacturer's instructions. PCR was performed using GoTaq Green 2x master mix (Promega), and results analysed using agarose gel electrophoresis. Band intensities were quantified using ImageJ. Alternatively, PCR was performed using PfU DNA Polymerase (Promega), and products were purified using AMPure XP magnetic beads (Beckman Coulter) following manufacturer's instructions. PCR products were barcoded, pooled and sequenced using a LSK109 library preparation kit on a single R.9.4.1 MinION flowcell (Oxford Nanopore Technologies) which was run for 4 hours. Raw FAST5 files were basecalled with Guppy 5 to produce raw FASTQ files. These files then underwent read correction using Canu 2.2[6] using the --correctReads parameter. Reads were aligned to the mm10 mouse reference genome using minimap2 (version 2.24[7]) using the parameters: *ax map-ont.* CRISPResso2 (https://crispresso.pinellolab.partners.org/submission)[8] was then run targeting the Rosa26 locus of the mm10 genome. Each read was assigned as an HR outcome if matching the template sequence, a TMEJ outcome if matching predicted TMEJ sequences, or an NHEJ outcome if containing non-TMEJ indels. TMEJ predictions were carried out using MEDJED (http://www.genesculpt.org/medjed/).

**Statistics and reproducibility.** All statistical tests used a two-sided unpaired *t*-test: N.S (not significant) p>0.05, "*" =p ≤0.05, "**" =p ≤ 0.01, "***" =p ≤ 0.001, "****" =p ≤ 0.0001. All experiments were repeated at least once and the number of biological replicates as are the numbers of event counted are reported for each experiment. To aid readability, statistics have only been shown between pertinent groups in figures.

### Antibodies.

***Antibody generation.*** Murine BRCA1 residues 1-300 bearing a Histidine tag and expressed in bacteria was used as an immunogen in Rabbits (Dundee cell products). The antibody is available on request to the corresponding authors subject to completion of a standard MTA.

| Antibody (clone) | Host | Supplier | Cat. number | Lot number | Technique | Conc | RRID |
|---|---|---|---|---|---|---|---|
| β-actin | Rabbit | Abcam | Ab8227 | GR3215935-1 | WB | 1:3000 | AB_2305186 |
| Human BARD1 | Rabbit | Abcam | ab226854 | GR3197067-8 | WB | 1:2000 | |

| | | | | | | | |
|---|---|---|---|---|---|---|---|
| Murine BARD1 | Rabbit | Gift from R.Baer (McCarthy et al., 2003) | N/A | N/A | WB/IF | 1:1000 | N/A |
| Human BRCA1 (D-9) | Mouse | Santa Cruz | Sc6954 | C2519 | IF | 1:500 | AB_626761 |
| Human BRCA1 (MS110) | Mouse | MERCK Millipore | OP94 | 3091924 | WB | 1:500 | AB_213438 |
| Murine BRCA1 (56E) | Rabbit | Gift from R.Baer | N/A | N/A | IF | 1:1000 | N/A |
| Murine BRCA1 (C40) | Rabbit | Dundee cell Products | Custom design | | IP | | |
| Murine BRCA1 (287.17) | Mouse | Santa Cruz | sc-135732 | D2121 | IF | 1:50 | AB_2243740 |
| Murine BRCA1 | Rabbit | Affinity Bioscience | AF6288 | 14f1430 | WB | 1:1000 | AB_2835138 |
| Pol theta | Rabbit | MyBiosource | MBS9612322 | 85i9616 | WB | 1:500 | |
| CldU (BrdU) | Rat | Abcam | Ab6326 | GR3173537-9 | Fibres/IF | 1:2000 | AB_305426 |
| Flag (M2) | Mouse | SigmaAldrich | F1804 | SLBT7654 | WB | 1:1000 | AB_262044 |
| IdU (BrdU) | Mouse | BD Biosciences | 347580 | 8151735 | Fibres | 1:500 | AB_400326 |
| γH2AX | Rabbit | Abcam | Ab2893 | GR3242597-1 | IF | 1:2000 | AB_303388 |
| PALB2 | Rabbit | Bethyl | A301.246A | | WB | 1:2000 | AB_890607 |
| RNF168 | Sheep | Novus Biologicals | AF7217 | | IF/WB | 1:1000 | AB_10971653 |
| RAD51 (Ab-1) | Rabbit | Calbiochem | PC130 | 3135376 | PLA | 1:100 | AB_2238184 |
| | | | | 3668125 | IF | 1:1000 | |
| Tubulin | Mouse | Santa Cruz | sc-5286 | H0613 | WB | 1:5000 | AB_628411 |
| Vinculin [EPR8185] | Rabbit | Abcam | Ab129002 | GR221671-50 | WB | 1:2000 | AB_11144129 |
| Donkey α Mouse AlexaFluor 488 | Donkey | Life technologies | A21202 | 1975519 | IF | 1:5000 | AB_141607 |
| Donkey α Rabbit AlexaFluor 488 | Donkey | Life technologies | A21206 | 1874771 | IF | 1:5000 | AB_2535792 |
| Donkey α Mouse AlexaFluor 555 | Donkey | Life technologies | A31570 | 1774719 | IF | 1:5000 | AB_2536180 |
| Donkey α Rabbit AlexaFluor 555 | Donkey | Life technologies | A31572 | 1945911 | IF | 1:5000 | AB_162543 |
| Donkey α Rat AlexaFluor 555 | Donkey | Life technologies | A21434 | 1987272 | IF | 1:5000 | AB_2535855 |
| Rabbit α Mouse HRP | Rabbit | Dako | P0161 | 20062080 | WB | 1:10000 | AB_2687969 |
| Swine α Rabbit HRP | Swine | Dako | P0217 | 20047666 | WB | 1:10000 | AB_2728719 |

**Supplementary Table 2** - siRNA sequences

| Target | siRNA Sequences | Supplier |
|---|---|---|
| NTC (Renilla | Sense: CUUACGCUGAGUACUUCGA[dT][dT] | Sigma |
| Luciferase) | Antisense: [Phos]UCGAAGUACUCAGCGUAA G[dT][dT] | Aldrich |
| mBRCA1 | Sense: GGAUUUAUCUGCCGUCCAA [dT][dT] | Sigma |
| | Antisense: [Phos] UUGGACGGCAGAUAAAUCC[ [dT][dT] | Aldrich |
| | Sense: GAACAGAGCAACUUGAAAC [dTdT] | Sigma |
| | Antisense: [Phos] AUUGUCUGUAUAGUCCACAGG [dT][dT] | Aldrich |
| mRNF168 | Sense: CCUUGGCUUCUCCUUUGAGUU [dT][dT] | Sigma |
| | Antisense: [Phos] AACUCAAAGGAGAAGCCAAGG [dT][dT] | Aldrich |
| mRAD52 | Sense: UUGAAGGUCAUCGGGUAAUUA [dT][dT] | Sigma |
| | Antisense: [Phos]UAAUUACCCGAUGACCUUCAA [dT][dT] | Aldrich |
| | Sense: ACUAUCUGAGGUCACUAAAUA [dT][dT] | |
| | Antisense: [Phos] UAUUUAGUGACCUCAGAUAGU [dT][dT] | |
| mPOL[illegible] | Sense: CCAGACUAAGAGUUCUCAUAA [dT][dT] | Sigma |
| | Antisense:[Phos] UUAUGAGAACUCUUAGUCUGG [dT][dT] | Aldrich |
| | Sense: CCAGGAAUCAAAGACGACAAU [dT][dT] | Sigma |
| | Antisense:[Phos] AUUGUCGUCUUUGAUUCCUGG [dT][dT] | Aldrich |
| | Sense: CACGGAAGAAAGCGUUGUUUA [dT][dT] | |
| | Antisense:[Phos] UAAACAACGCUUUCUUCCGUG [dT][dT] | |

| | | |
|---|---|---|
| mRADX | Sense: CAUAGAGGCCAGCCGUAUA [dT][dT]<br>Antisense:[Phos] UAUACGGCUGGCCUCUAUG [dT][dT]<br>Sense: GAAAGUAUUCCACGGAAAUUU [dT][dT]<br>Antisense:[Phos] AAAUUUCCGUGGAAUACUUUC [dT][dT]<br>Sense: GGGAUAAUUACUGCUAUAAAG [dT][dT]<br>Antisense:[Phos] CUUUAUAGCAGUAAUUAUCCC [dT][dT] | Sigma Aldrich |
| hBRCA1 | Sense: GCUCCUCUCACUCUUCAGU[dTdT]<br>Antisense: [Phos]ACUGAAGAGUGAGAGGAGC[dT][dT]<br><br>Sense: AAGCUCCUCUCACUCUUCAGU[dT][dT]<br>Antisense: [Phos]ACUGAAGAGUGAGAGGAGCUU[dT][dT] | Sigma Aldrich |

**Supplementary Table 3** – primer, template, and gRNA sequences

| Gene | Point mutation (human) | Point mutation (murine) | Sequences (5'-3-) | Supplier |
|---|---|---|---|---|
| BARD1 | L44R | L38R | Forward: GCTTGCCCGCCGGGAGAAGCTGCTG<br>Reverse: ACTGGGCATCCTGAGCCAACACAG | Sigma Aldrich |
| | F133A/D135A/A136E<br>(AAE) | F125A/D127A/A128E<br>(AAE) | Forward: CATTTTTATTGAATTCTTCTTCCTTTCTTCAGCACCAGCTAAA CTTGCCCTAGATG TGTTGTCTTTTGAAT<br>Reverse: ATTCAAAAGACAACACATCTAGGGCAAGTTTAGCTGGTGCT GAAGAAAGGAAGAAGAATTCAATAAAAATG | Sigma Aldrich |
| | A460T | A448T | Forward: CGGTGTCCATCCAGTATGGTCTTTAACATTTGGGT<br>Reverse: ACCCAAATGTTAAAGACCATACTGGATGGACACCG | Sigma Aldrich |
| | D712A | D700A | Forward: GATGGTCTGAGTCACAGCACTGTCTGGCTTGGG<br>Reverse: CCCAAGCCAGACAGTGCTGTGACTCAGACCATC | Sigma Aldrich |
| BRCA1 | M18T | - | Forward: CAAAATGTCATTAATGCTTTGCAGAAAATCTTAGAGTG | Sigma Aldrich |
| Rosa 26 F | | | AGAAAACTGGCCCTTGCCATT | Sigma Aldrich |
| Roas26 R | | | CAGCCTCGATTTGTGGTGTATG | Sigma Aldrich |
| Rosa26 HR assay template | | | GGGGGAGTCGTTTTACCCGCCGCCGGCCGGGCCTCGTCGTCTGATTGGC TCTCGGGGCCCAGAAAACTGGCCCTT GCCATTGGCTCGTGTTCGTGCAA GTTGAGTCCATCCGCCGGCCAGCGGGGGCGGCGAGGAGGCGCTCCCAG GTTC CGGCCCTCCCCTCGGCCCCGCGCCGCAGAGTCTGGCCGCGCGCCC CTGCGCAACGTGGCAGGAAGCGCGCGCTG GGGGCGGGGACGGGCAGT AGGGCTGAGCGGCTGCGGGGCGGGTGCAAGCACGTTTCCGACTTGAGT TGCCTCAA GAGGGGCGTGCTGAGCCAGACCTCCATCGCGCACTCCGGG GAGTGGAGGGAAGGAGCGAGGGCTCAGTTGGGCT GTTTTGGAGGCAG GAAGCACTTGCTCTCCCAAAGTCGCTCTGAGTTGTTATCAGTAAGGGAGC TGCAGTGGAGTAG GCGGGGAGAAGGCCGCACCCTTCTCCGGAGGGGG GAGGGGAGTGTTGCAATACCTTTCTGGGAGTTCTCTGCTGC CTCCTGGC TTCTGAGGACCGCCCTGGGCCTGGGAGAATCCCTTCCCCCTCTTCCCTCG TGATCTGCAACTCCAGTCTTTCTAGAAGtactGCGGGAGTCTTCTGGGCA GGCTTAAAGGCTAACCTGGTGTGTGGGCGTTGTCCTGCAGGGG AATTG AACAGGTGTAAAATTGGAGGGACAAGACTTCCCACAGATTTTCGGTTTT GTCGGGAAGTTTTTTAATAGGGG CAAATAAGGAAAATGGGAGGATAG GTAGTCATCTGGGGTTTTATGCAGCAAAACTACAGGTTATTATTGCTTGT GAT CCGCCTCGGAGTATTTTCCATCGAGGTAGATTAAAGACATGCTCAC CCGAGTTTTATACTCTCCTGCTTGAGATCC TTACTACAGTATGAAATTAC | Genscript |

| | |
|---|---|
| | AGTGTCGCGAGTTAGACTATGTAAGCAGAATTTTAATCATTTTTAAAGA GCCCAGTAC TTCATATCCATTTCTCCCGCTCCTTCTGCAGCCTTATCAAAA GGTATTTTAGAACACTCATTTTAGCCCCATTTTCATT TATTATACTGGCTT ATCCAACCCCTAGACAGAGCATTGGCATTTTCCCTTTCCTGATCTTAGAA GTCTGATGACTCATGAAACCAGACA |
| Rosa26 gRNA | ACTCCAGTCTTTCTAGAAGA |

**Supplementary Table 4** - Details of DNA damaging agents and inhibitors

| Inhibitor | Company | Catalogue Number | Target | Concentration |
|---|---|---|---|---|
| Olaparib | Selleckchem | S1060 | PARP | 1-20 µM |
| Aphidicolin | Sigma-Aldrich | A0781 | DNA polymerase α | 5-100 nM |
| Cisplatin | Sigma-Aldrich | PHR1624 | DNA-crosslinker | 1-5 µM |
| CldU | Sigma-Aldrich | C6891 | Thymidine analogue | 250 µM |
| EdU | Thermofisher | A10044 | Thymidine analogue | 10 µM |
| IdU | Sigma-Aldrich | I7125 | Thymidine analogue | 25 µM |
| Hydroxyurea | Sigma-Aldrich | H8627 | Ribonucleoside reductase | 1-10 mM |
| Colcemid | Gibco | 15212012 | Microtubule poison | 0.05 µg/ml |
| 6-OH-L-Dopamine | Sigma | H2380 | RAD52 | 0.15-5 µM |
| DI03 | Sigma | SML2496 | RAD52 | 0.5-10 µM |
| Novobiocin | Selleckchem | NSC2382 | Polymerase Theta | 50-200 µM |
| ART558 | Artios | n/a | Polymerase Theta | 2.5-20 µM |

| | | | | |
|---|---|---|---|---|
| L67 | Sigma-Aldrich | SML1797 | DNA Ligase I/III | 1-2 µM |
